# Functional Analysis of the Zinc Finger Modules of the *S. cerevisiae* Splicing Factor Luc7

**DOI:** 10.1101/2024.02.04.578419

**Authors:** Tucker J. Carrocci, Samuel DeMario, Kevin He, Natalie J. Zeps, Cade T. Harkner, Guillaume Chanfreau, Aaron A. Hoskins

## Abstract

Identification of splice sites is a critical step in pre-mRNA splicing since definition of the exon/intron boundaries controls what nucleotides are incorporated into mature mRNAs. The intron boundary with the upstream exon is initially identified through interactions with the U1 snRNP. This involves both base pairing between the U1 snRNA and the pre-mRNA as well as snRNP proteins interacting with the 5’ splice site/snRNA duplex. In yeast, this duplex is buttressed by two conserved protein factors, Yhc1 and Luc7. Luc7 has three human paralogs (LUC7L, LUC7L2, and LUC7L3) which play roles in alternative splicing. What domains of these paralogs promote splicing at particular sites is not yet clear. Here, we humanized the zinc finger domains of the yeast Luc7 protein in order to understand their roles in splice site selection using reporter assays, transcriptome analysis, and genetic interactions. While we were unable to determine a function for the first zinc finger domain, humanization of the second zinc finger domain to mirror that found in LUC7L or LUC7L2 resulted in altered usage of nonconsensus 5’ splice sites. In contrast, the corresponding zinc finger domain of LUC7L3 could not support yeast viability. Further, humanization of Luc7 can suppress mutation of the ATPase Prp28, which is involved in U1 release and exchange for U6 at the 5’ splice site. Our work reveals a role for the second zinc finger of Luc7 in splice site selection and suggests that different zinc finger domains may have different ATPase requirements for release by Prp28.

## INTRODUCTION

Eukaryotic pre-messenger RNA (pre-mRNA) is modified to remove internal sequences called introns by pre-mRNA splicing. Splicing is a highly conserved process across eukaryotes, and it is carried out by the large macromolecular machine known as the spliceosome. Spliceosomes are composed of five U-rich small nuclear ribonucleoprotein (snRNP) complexes (U1, U2, U4, U5, U6) and dozens of auxiliary proteins (Wilkinson et al. 2020). These factors assemble *de novo* on every intron through the recognition of splice sites in order to form active spliceosomes and catalyze intron removal. Defects in this process often lead to aberrantly spliced RNA products and can be causative for human disease (Love et al. 2023).

Splicing often begins with base pairing of the 5’ end of the U1 snRNA to the intron 5’ splice site (5’ss) at the exon-intron boundary to form the spliceosome E complex (Ruby and Abelson 1988; Shcherbakova et al. 2013; Fica 2020). U1 remains associated with the 5’ss during spliceosome assembly but must be released by the ATPase Prp28 during the pre-B to B complex transition to allow for U6 snRNA pairing to the 5’ss (Staley and Guthrie 1999). The sequence of the 5’ss is highly conserved in *Saccharomyces cerevisiae* (hereafter yeast) whereas human 5’ss are more divergent (Spingola et al. 1999; Roca et al. 2013). In addition, human splicing often requires auxiliary splicing factors to direct U1 recruitment and stabilize U1 on the pre-mRNA (Matlin et al. 2005; Espinosa et al. 2022). In addition to RNA base pairing, the U1 snRNA:5’ss duplex is also stabilized by protein components of the U1 snRNP. In yeast, the U1 snRNP proteins Yhc1 and Luc7 flank the duplex (**Fig. 1a**). These proteins are conserved in humans; however, the human homolog of Yhc1 (U1-C) is considered to be a core component of the U1 snRNP while the human homologs of Luc7 are auxiliary splicing factors. Cryo-electron microscopy has revealed the architecture of both isolated yeast U1 snRNP and of the spliceosome A complex (which contains the U1 and U2 snRNPs) and indicated that Yhc1 and Luc7 interact with U1 snRNA:5’ss duplex through their zinc finger (ZnF) domains (Li et al. 2017; Plaschka et al. 2018; Li et al. 2019). Alterations in the Yhc1 and Luc7 ZnF domains have been shown to impair splicing *in vivo* and can be lethal (Schwer and Shuman 2014; Agarwal et al. 2016).

**Figure 1.**
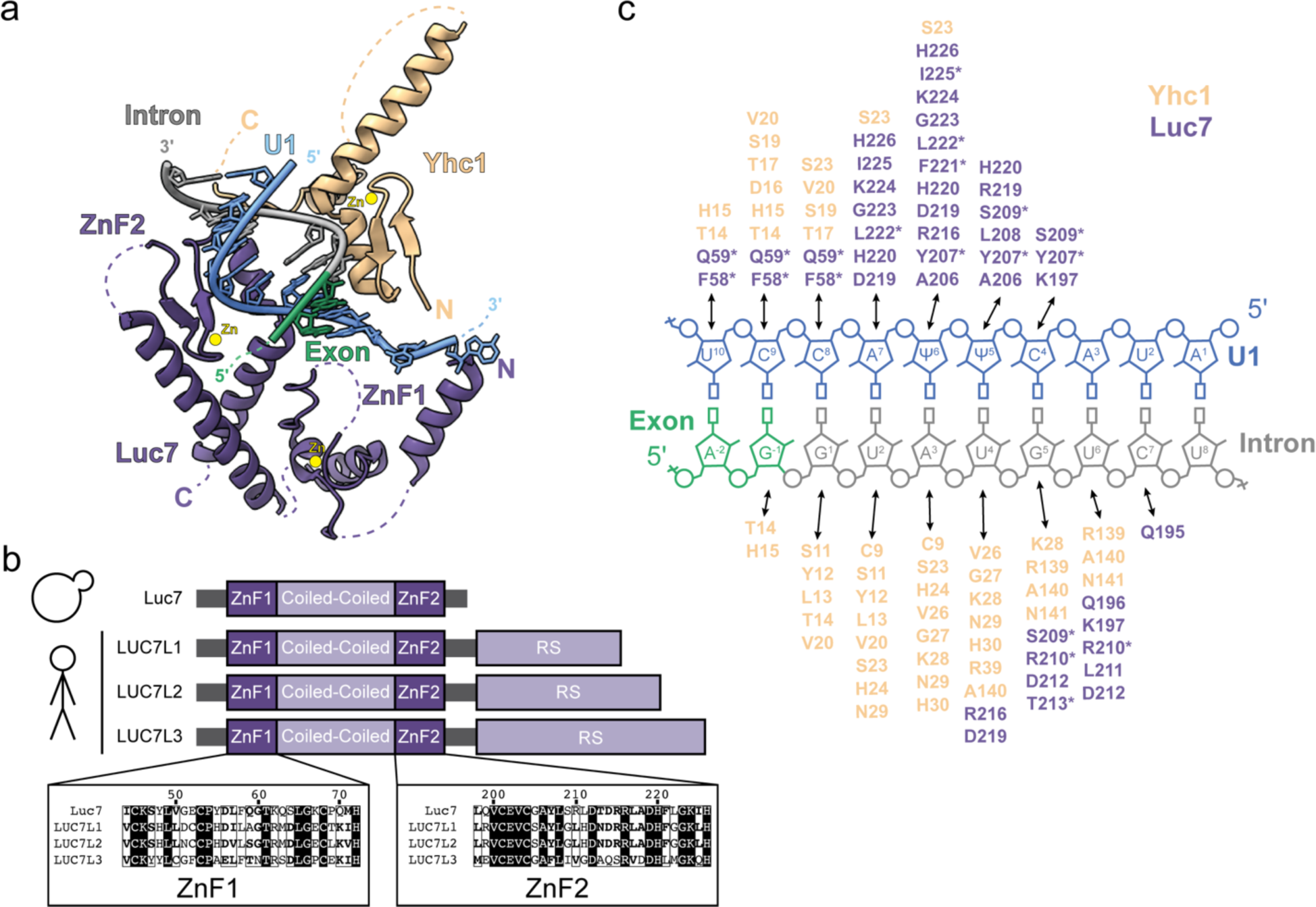
The U1:5’ss duplex contacts Yhc1/U1-C and Luc7/LUC7L. (**a)** U1:5’ss base pairing is stabilized by the Yhc1 and Luc7 ZnF domains in the structure of the yeast pre-spliceosome (PDB 6G90). Luc7 ZnF2 contacts the 5’ss duplex whereas ZnF1 lies in the direction of exon 1, which are both partially unresolved. (**b**) Schematic showing *LUC7* and the three human paralogs. A sequence alignment shows similarity of the ZnF domains. (**c**) Schematic showing Yhc1 (beige) and Luc7 (purple) residues located within 6 Å of the U1:5’ss duplex (blue/grey). Asterisks indicate residues that are not conserved between yeast and human Luc7 proteins.

Luc7 is highly conserved among eukaryotes, and vertebrates have three Luc7 paralogs: LUC7L, LUC7L2, and LUC7L3 (Fortes et al. 1999) (**Fig. 1b**). Each paralog is proposed to control a distinct subset of alternative splicing events (Daniels et al. 2021; Jourdain et al. 2021). Loss of LUC7L2 has been implicated in myelodysplastic syndromes and related neoplasms and can lead to changes in glycolysis and metabolic reprogramming (Daniels et al. 2021; Jourdain et al. 2021). Specificities of the LUC7L paralogs for different 5’ss have been proposed to be due to differential recruitment to target RNAs and/or different sequence preferences (Daniels et al. 2021; Kenny et al. 2022). Understanding the molecular basis for specificity and function of the LUC7L proteins is important to establish how their loss can lead to disease or changes in metabolism.

Here, we examine differences among human LUC7L paralogs using a yeast model system. We generated yeast and human chimeric Luc7 proteins, focusing on the ZnF domains (ZnF1 and ZnF2), and assayed them for differences in splicing and splice site usage. We were unable to detect changes in splicing of a reporter pre-mRNA due to changes in Luc7 ZnF1. In contrast, we find that humanization of yeast Luc7 ZnF2 to mirror that in human LUC7L and LUC7L2 improves growth of yeast expressing splicing reporters containing non-consensus 5’ss, but yeast are inviable if ZnF2 is humanized to resemble LUC7L3 ZnF2. Consistent with reporter substrate data, we identify a subset of yeast transcripts that show altered processing upon humanization of Luc7 *in vivo*. Finally, we show that humanized Luc7 does not bypass the requirement for Prp28 in splicing but does suppress cold-sensitive (*cs*) phenotypes observed with Prp28 mutants. Taken together, these data show that the Luc7 ZnF2 domain facilitates 5’ss selection.

## RESULTS

### Humanized Luc7 ZnF Proteins Support Yeast Viability Except for LUC7L3 ZnF2

To investigate the human Luc7 homologs, we generated a yeast “shuffle” system to introduce Luc7 mutants in yeast. We deleted the chromosomal *LUC7* gene while maintaining yeast viability by expression of wild type (WT) Luc7 from a low-copy, *URA3/CEN6*-containing plasmid. Plasmid expression of Luc7 had no effect on yeast viability in standard growth conditions (**Fig. S1A**). To introduce human Luc7 homologs in yeast, we inserted yeast codon-optimized gene fragments encoding truncations of the open reading frames of LUC7L, LUC7L2, or LUC7L3 into a low-copy, *TRP1/CEN6*-containing plasmid containing the native *LUC7* promoter and terminator sequences. These truncations coded for the predicted structured regions of the proteins but did not include the C-terminal RS domains. Each of the constructs also included a 3×HA epitope tag at the protein C-terminus.

After transformation of yeast with these plasmids and subsequent 5-FOA selection, we found that none of the human homologs were able to support yeast growth as the sole copy of Luc7 (**Fig. S1B**). We confirmed expression of the proteins by western blotting. While each protein was expressed, LUC7L and LUC7L2 were less abundant than either Luc7 or LUC7L3 (**Fig. S1C**). In addition, we were unable to rescue viability when these proteins were expressed under control of a strong *TDH3* promoter (OE, **Fig. S1B**). The inability of the human LUC7L proteins to fully compensate for yeast Luc7L has also been recently reported by Chalivendra *et al (Chalivendra et al. 2023)*. Based on these results, we instead focused on targeted mutations to humanize Luc7.

We analyzed the cryo-EM structure of the yeast spliceosome A complex to identify Luc7 amino acids located within 6 Å of the U1 snRNA/5’ss duplex (**Fig. 1a**, **c**) (Plaschka et al. 2018). The amino acids predominantly come from ZnF2; however, we also decided to study ZnF1 (which is likely located further from the RNA duplex) since previous work showed genetic interactions between ZnF1 mutations and a number of splicing factors (Agarwal et al. 2016). We identified the corresponding amino acids in ZnF1 and ZnF2 of the human LUC7L paralogs based on a multiple sequence alignment generated by Clustal Omega (Sievers and Higgins 2014) (**Fig. 1b**) and humanized yeast Luc7 accordingly by site-directed mutagenesis of the non-conserved amino acids (**Table S1**). We then expressed the humanized, C-terminally 3×HA-tagged Luc7 variants in yeast on plasmids and under control of the native Luc7 promoter. We refer to mutants of ZnF1 as ZnF1_L, L2, or L3 corresponding to ZnF1 from LUC7L, LUC7L2, or LUC7L3. The mutations needed to humanize ZnF2 for LUC7L and LUC7L2 are identical. We refer to mutants of ZnF2 as ZnF2_L/L2 or L3.

Luc7 chimeric proteins containing one or both ZnFs from LUC7L or LUC7L2 were viable (**Fig. 2**). While a Luc7 chimera of ZnF1_L3 was also viable, a chimera containing ZnF2_L3 was not even though it was well-expressed in yeast (**Fig. S1d**). ZnF2_L3 did not support viability even if ZnF1 was also humanized to that from LUC7L3 or any other LUC7L paralog. From these experiments we conclude that while most ZnF modules from human Luc7 homologs can support yeast splicing, ZnF2_L3 is not functionally equivalent to either ZnF2 from Luc7 or LUC7L/L2.

**Figure 2.**
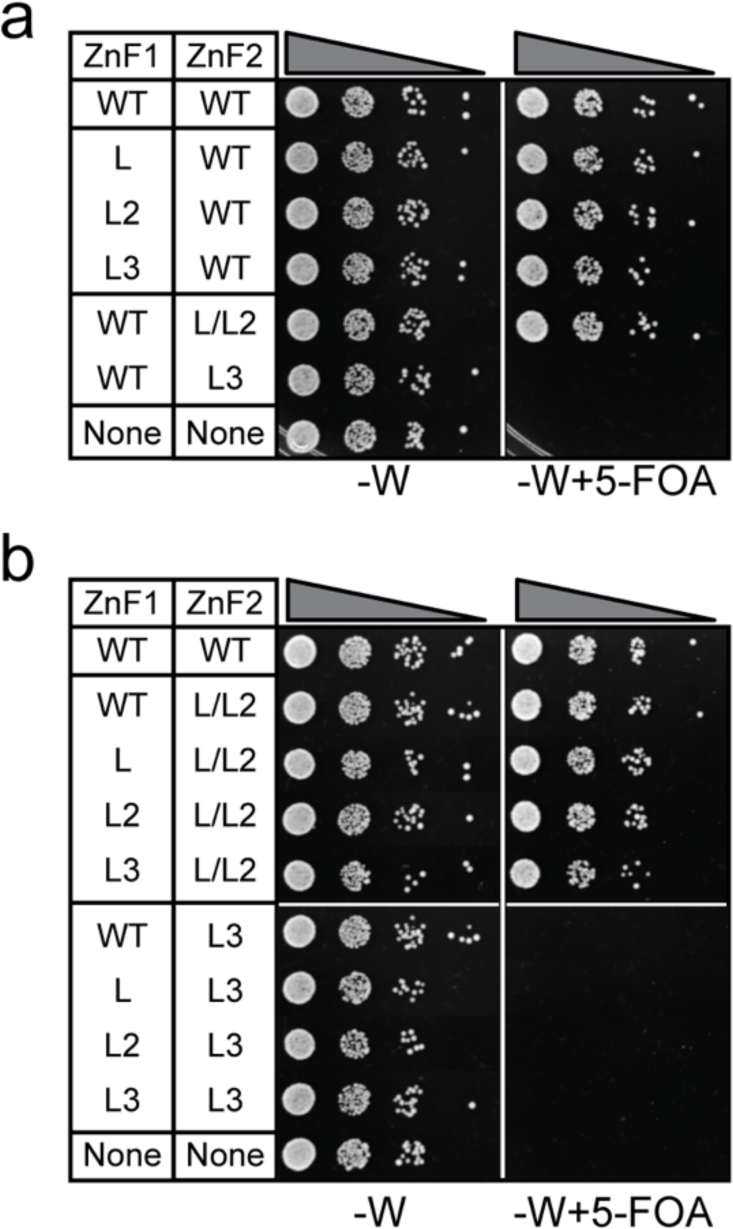
Yeast are viable with humanized Luc7 ZnF1 and ZnF2. (**a**) Luc7 ZnF1 mutants are viable in yeast. The ZnF2_L/L2 mutant is also viable whereas ZnF2_L3 is lethal. (**b**) Expressing Luc7 with mutations in both ZnF1 and ZnF2 does not rescue the lethality associated with ZnF2_L3.

### ZnF2 but not ZnF1 Changes 5’ss Usage of a Yeast Splicing Reporter

We next used the ACT1-CUP1 reporter assay to probe how humanization of the Luc7 ZnF domains influences 5’ss selection. In this assay, yeast growth in the presence of various concentrations of Cu^2+^ is correlated with splicing of the reporter and mRNA production (**Fig. 3a**) (Lesser and Guthrie 1993a). In the case of ZnF1, we used ACT1-CUP1 reporters with substitutions from −3 to −1 on the exonic side of the 5’ss since this region is closest to the binding site of ZnF1 (**Fig. 1**, **3a**). The WT ACT1-CUP1 substrate normally only contains a single potential base pair with the U1 snRNA within the −3 to −1 region (−1G could pair with U1 snRNA C9, **Fig. 1c**). G-1A and G-1C substitutions in the ACT1-CUP1 should therefore disrupt all pairing within this region, while a U-2A or C-3A should strengthen the interaction by allowing pairing with either U10 or U11. To increase sensitivity of the assay, we also incorporated mutations at the +2 (U2A) or +5 (G5A) positions since substitutions at −3 to −1 have no discernible effect on the splicing of a substrate with a consensus 5’ss.

**Figure 3.**
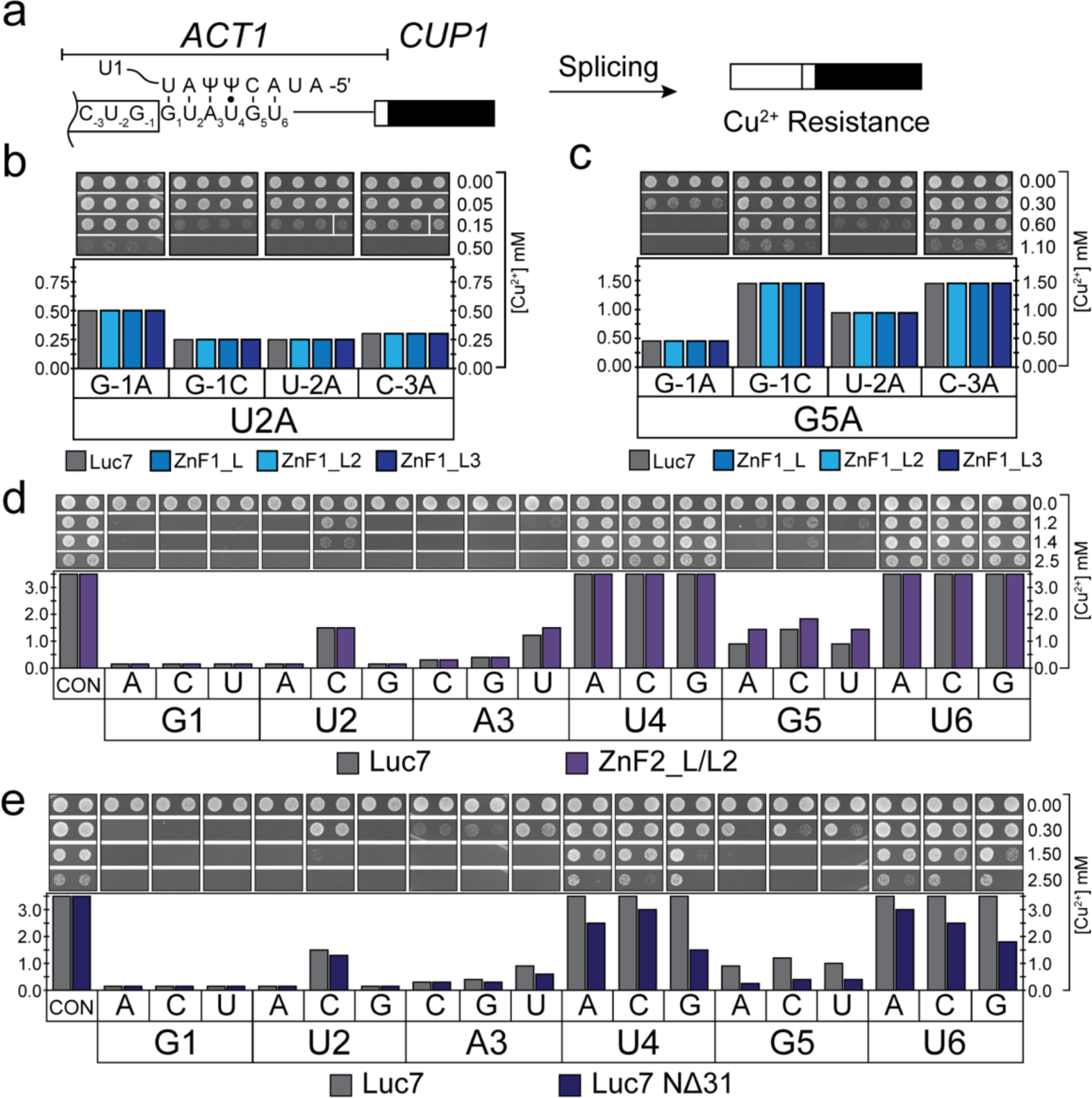
Mutation of the ZnF domains of Luc7 can have specific impacts on 5’ss usage. (**a**) Schematic representation of the *ACT1-CUP1* reporter pre-mRNA and splicing assay. Proper processing of the splicing reporter confers resistance to Cu^2+^ in the growth media. (**b**) *ACT1-CUP1* assays using a reporter with substitutions in the 5’ss sequence U2A (G<u>a</u>AUGU) and the indicated changes in the exon. ZnF1 mutants have no obvious effect on splicing. (**c**) Same as in **b** but using a reporter with the 5’ss G5A (GUAU<u>a</u>U). (**d**) ACT1-CUP1 assays using ZnF2_L/L2 to assay intronic positions of the 5’ss. Mutation of Luc7 ZnF2 improves splicing of A3U, G5A, G5C and G5U but does not affect other reporters. (**e**) ACT1-CUP1 assays using the NΔ31 truncation mutant of Luc7. Shown is a representative ACT1-CUP1 assay from three experimental replicates.

None of the ZnF1 chimeras changed yeast tolerance to Cu^2+^ for any of the ACT1-CUP1 reporters harboring substitutions at −3 to −1 (**Fig. 3b**, **c**). No changes were observed even when ZnF2 was also modified to that from LUC7L/L2 (**Fig. S2**). As expected, we also did not observe changes in splicing due to ZnF1 substitutions with reporters carrying substitutions at the +1 to +6 positions, likely because these substitutions are located distal to the ZnF1 interaction site (**Fig. S3**). From these results, we conclude that paralog-specific contacts of ZnF1 made by these humanized Luc7 variants are unlikely to significantly influence splicing of the ACT1-CUP1 reporter in this assay.

For studies of ZnF2, we focused on using ACT1-CUP1 reporters harboring substitutions within the intronic portion of the 5’ss (+1 to +6) since ZnF2 contacts this region (**Fig. 1c**). When ZnF2 was replaced by ZnF2_L/L2, we observed increased Cu^2+^ tolerance of yeast with ACT1-CUP1 reporters containing substitutions at the +3 (A3U) and +5 positions (G5A, C, U) (**Fig. 3d**), in agreement with contacts between Luc7 and the snRNA/5’ss duplex. Since the high levels of Cu^2+^ tolerance observed with some reporters precluded analysis (*i.e.,* U4 and U6 substitutions), we also carried out assays in which a second mutation was incorporated at the +2 position (U2A). We were unable to find any effect of ZnF2_L/L2 with these doubly substituted reporters for changes at the +4 or +6 positions (**Fig. S4**). These results suggest that a function of ZnF2 in human LUC7L and LUC7L2 could be to promote splicing at weak 5’ss containing mismatches to the U1 snRNA at the +3 and +5 positions of the 5’ss.

### Deletion of a Luc7/Sm Ring Interaction Decreases Usage of Nonconsensus Splice Sites

We wondered if other Luc7 mutants, not confined to the ZnF domains, would also show changes in splicing at +3 and +5. The Luc7 NΔ31 mutation abolishes contact between Luc7 and the U1 snRNP Sm protein ring and was previously shown to impair splicing of some pre-mRNAs, cause synthetic lethality with a number of other mutations, and bypass the need for Prp28 (Agarwal et al. 2016). Since this mutant was previously well-characterized, we carried out ACT1-CUP1 assays with Luc7 NΔ31-expressing yeast.

Unlike the ZnF2_L/L2 substitution, Luc7 NΔ31 caused a loss of yeast tolerance to Cu^2+^ using reporters with multiple substitutions at the +2 to +6 positions (**Figs. 3e**, **S4**). The only position that was unaffected was +1. This suggests that pairing at this position and enforcing selection of the nearly invariant +1G at the 5’ss may be due to other snRNP components, likely Yhc1/U1-C which makes extensive contacts with the duplex at this position (**Fig. 1c**) (Kondo et al. 2015; Hansen et al. 2022). Surprisingly, Luc7 NΔ31 even decreased Cu^2+^ tolerance in yeast with U4A and U4G substitutions which should result in increased pairing to Ψ5 in the snRNA. This would seem to indicate that WT Luc7 has evolved to function preferentially on substrates containing mismatches (*i.e.,* a U) at the +4 position of the 5’ss. Interestingly, the +4 nucleotide is ultimately juxtaposed with A49 in the U6 snRNA during splicing catalysis, and +4U could form a canonical base pair with A49 (Kandels-Lewis and Seraphin 1993; Lesser and Guthrie 1993b). Luc7 may help the snRNP to tolerate a 5’ss/U1 snRNA mismatch that ultimately increases 5’ss base pairing to U6.

### Luc7 Variants Perturb Splicing of Endogenous Yeast pre-mRNAs

5’ss sequences found in endogenous yeast introns are less diverse than the reporters we used in the ACT1-CUP1 assays. For example, we do not believe there are any naturally-used 5’ss with an adenosine located at the +5 position even though we observed increased Cu^2+^ tolerance with the G5A reporter and ZnF2_L/L2. We first tested two endogenous transcripts, *SUS1* and *RPL22B*, for changes in splicing due to Luc7 ZnF2_L/L2 or Luc7 NΔ31. We chose these transcripts because changes in *SUS1* splicing had previously been reported for Luc7 NΔ31 (Agarwal et al. 2016) and *RPL22B* has a cryptic, intronic 5’ss (GUUUGU) with an A3U substitution relative to the consensus (Kawashima et al. 2014). Since the splicing of ACT1-CUP1 reporters with A3U is stimulated by Luc7 ZnF2_L/L2 (**Fig. 3d**), we wondered if the humanized protein would also stimulate use of the cryptic site in *RPL22B*.

We analyzed splicing of *SUS1* and *RPL22B* by RT-PCR using yeast strains expressing Luc7-ZnF2_L/L2 or Luc7 NΔ31. UPF1 was also deleted in these strains to block nonsense mediated decay (NMD) and facilitate detection of *RPL22B* isoforms (Kawashima et al. 2014). Both Luc7 variants somewhat inhibited splicing of both *SUS1* and *RPL22B* with Luc7 NΔ31 showing a greater splicing defect (**Fig. S5**). We did not observe a substantial increase in use of the cryptic 5’ss in *RPL22B* due to Luc7 ZnF2_L/L2. However, Luc7 NΔ31 accumulated unspliced *RPL22B* and had comparatively less usage of the cryptic 5’ss than either WT Luc7 or Luc7 ZnF2_L/L2. This is consistent with the ACT1-CUP1 assays and with Luc7 NΔ31 inhibiting splicing of weak, nonconsensus 5’ss including A3U (**Fig. 3e**).

To further probe the effects of the Luc7 variants on the yeast transcriptome, we used total RNA-seq to analyze changes in splicing efficiency genome-wide. RNA sequencing was performed in a *upf1*Δ genetic background deficient for nonsense-mediated mRNA decay (NMD). The *upf1*Δ genetic background not only helps to stabilize unspliced mRNAs harboring premature termination codons but also facilitates a more direct analysis of splicing efficiency without confounding effects of differential isoforms stability (Sayani et al. 2008; Kawashima et al. 2014).

mRNA expression was globally unchanged when Luc7 ZnF2_L/L2 was expressed compared to WT, as no genes showed significant differential expression (*p* value < 1e-5). The Luc7 NΔ31 strain had 9 differentially expressed genes (**Fig. S6A**): *YBL059W*, ATP5, IMD2, DGR2, YLR366W, RPS22B*, YML131W, HRB1*, LYS9* (* indicates an intron-containing gene). *HRB1* is interesting due to its reported role in selective export of spliced mRNAs (Hackmann et al. 2014). Its modest upregulation (∼1.6 fold) is likely due to the accumulation of unspliced transcripts. It is worth noting that analysis of *HRB1* RNAs also showed a significant fraction of unspliced reads in the control *upf1*Δ strain expressing WT Luc7 (approximately 1:1), indicative of suboptimal splicing even in the presence of the WT protein.

To calculate splicing efficiency, we quantified reads which could be unambiguously assigned as either spliced junctions or unspliced reads (*i.e.*, spanning both exonic and intronic regions). For each intron annotation we first calculated the ratio of unspliced reads to spliced reads. We then compared these ratios for each mutant relative to the WT control. Both Luc7-ZnF2_L/L2 and Luc7 NΔ31 had a negative impact on global splicing efficiency compared to WT (**Fig. 4A**). However, the effect was not universal as both mutants showed increased splicing efficiency of a select few transcripts. We grouped each intron by its change in splicing efficiency in each mutant compared to WT and analyzed splicing sequences for each group by generating sequence logos of the 5’ss, branch points, and 3’ss (**Figs. 4A**, **S6B**).

**Figure 4.**
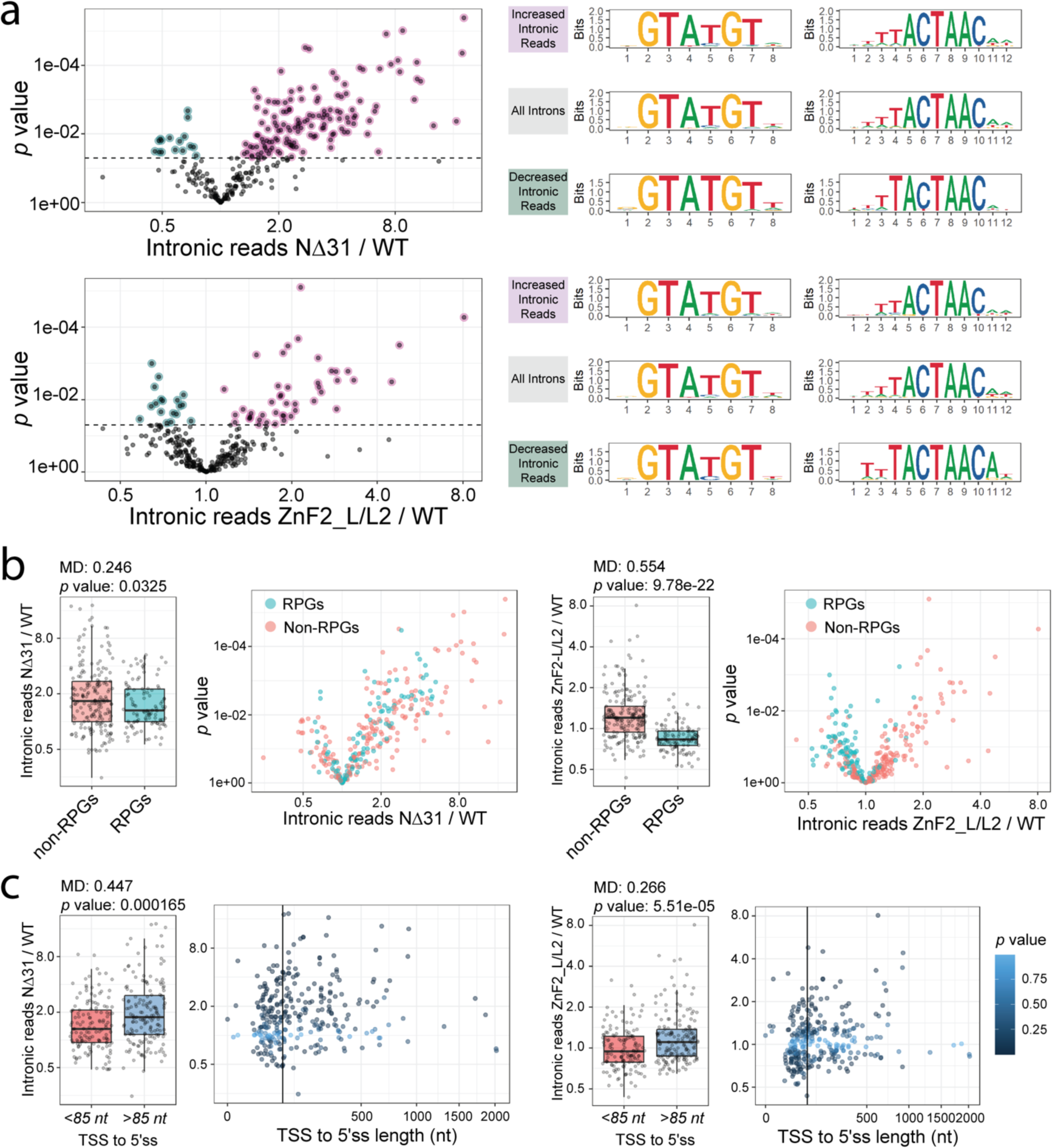
Analysis of RNA-seq data from cells expressing WT, NΔ31, or ZnF2_L/L2 Luc7 Proteins. (**a**) Plots showing changes in splicing efficiencies for each intron in Luc7 NΔ31 (Top) and Luc7 ZnF2_L/L2 (Bottom). *p* values are the results of unequal variances t-tests using geometric means between 3 replicates. Introns with statistically significant changes in splicing efficiency were categorized into increased or decreased intronic reads in mutants over WT, *p* value < 0.05 (dashed line). Sequence logos for 5’ splice sites (middle) and branch points (right) are shown for each category. (**b**) Changes in splicing efficiencies in Luc7 NΔ31 (Left) and Luc7 ZnF2_L/L2 (Right) vs WT for ribosomal protein genes. Cyan points indicate ribosomal protein genes. Histogram *p* values are the results of unequal variances t-tests using geometric means. (**c**) Plots showing transcriptional start site to 5’ss distances vs changes in splicing efficiency for Luc7 NΔ31 (Left) and Luc7 ZnF2_L/L2 (Right) vs WT Luc7. Color indicates *p* values. Black vertical line indicates 85 nt. (MD - Difference of geometric means)

We found that introns with increased splicing efficiency with Luc7 NΔ31 had near perfect consensus 5’ss (GUAUGU). In line the ACT1-CUP1 data, introns with non-“GUAUGU” 5’ss had an average of 45% more unspliced reads with Luc7 NΔ31 compared to WT (**Fig. S6C**). The 5’ss sequence had surprisingly little effect with the Luc7 ZnF2_L/L2. Only one of the common, non-consensus 5’ss showed a statistically significant change in splicing efficiency: introns with a GUACGU 5’ss had 18% fewer unspliced reads on average (**Fig. S6C**). This is consistent with the position of the ZnF2 region in contacting the U1 snRNA/5’ss duplex near the U4 position, but we did not observe a similar change at U4 substitutions in our ACT1-CUP1 assays (**Figs. 3d, S4**).

The introns with increased splicing efficiency in both the Luc7 ZnF2_L/L2 and NΔ31-expressing strains showed a strong enrichment for the consensus U at position one of the branch point consensus (<u>U</u>ACUAAC, underlined) as well as a more modest enrichment for the consensus C at position 7 (UACUAA<u>C</u>). Interestingly, the introns with increased splicing efficiency using the Luc7 ZnF2_L/L2 variant showed an enrichment for an A immediately downstream of the branch point (UACUAAC<u>A</u>). An adenosine at this position would facilitate formation of an additional base pair with the U2 snRNA at U2 nucleotide U33. Base pairing between the U2 snRNA and intron is seen at this position in cryo-EM structures of the U2 snRNP (Plaschka et al. 2018; Zhang et al. 2024). This suggests that increased base pairing between U2 and the branch point may compensate for the deleterious effects of the ZnF2 L/L2 variant.

We also found that the strain expressing Luc7 ZnF2_L/L2 was able to splice pre-mRNAs of ribosomal protein genes (RPGs) protein pre-mRNAs with much higher efficiency than pre-mRNAs of non-RPGs, while the strain expressing Luc7 NΔ31 showed little difference between the two (**Fig. 4B**). In an attempt to explain this observation, we checked for correlations between intron length and splicing efficiency as well as between the transcription start site (TSS) to 5’ss distance and splicing efficiency, since introns found in RPGs are typically longer than those of non-RPGs (Spingola et al. 1999). While both were statistically significant with the Luc7 ZnF2_L/L2 mutant, neither was a better explanatory variable for splicing efficiency than ribosomal vs non-ribosomal protein gene (**Figs. 4C**, **S6D**).

While splicing efficiency in the Luc7 NΔ31-expressing strain was not significantly correlated with ribosomal vs non-ribosomal protein gene or intron length, it did show a significant correlation with the distance from TSS to the 5’ss (**Fig. 4C**) with longer distances from the TSS corresponding to generally lower splicing efficiency. Since our data (**Figs. 3e**, **4**) and that by Agarwal and coworkers (Agarwal et al. 2016) suggest that Luc7 NΔ31 is less able to stabilize the U1 snRNA/5’ss interaction, it would seem to indicate that this becomes even worse the further from the TSS and (possibly) the 5’ end of the nascent transcript. We do not know exactly what factors are responsible for this behavior, but Luc7 NΔ31 is synthetically lethal with deletion of the nuclear cap binding complex (CBC) (Agarwal et al. 2016). It is possible that the ability of the CBC to stabilize U1 on the pre-mRNA (Gornemann et al. 2005; Larson and Hoskins 2017) and compensate for Luc7 NΔ31 is less effective at greater TSS to 5’ss distances. Similar RNA cap to 5’ss distance dependence for the function of Luc7 has been remarked upon previously with the presence of Luc7 favoring selection of cap-proximal 5’ss (Puig et al. 2007).

### Luc7 ZnF2_L/L2 Alters the Requirement for Prp28 During Splicing

We next assayed the Luc7 mutants for an altered requirement of the DEAD-box ATPase Prp28 for U1/U6 base pairing exchange at the 5’ss during spliceosome activation (**Fig. 5A**) (Staley and Guthrie 1999). It has previously been shown that Luc7 NΔ31 can completely bypass the need for Prp28 activity and that the *PRP28* gene is no longer essential when Luc7 NΔ31 is present (Agarwal et al. 2016). Unlike Luc7 NΔ31, Luc7 ZnF2_L/L2 does not bypass the requirement for the presence of Prp28 during splicing (**Fig. 5B**). We then tested whether Luc7 ZnF2_L/L2 could alter the demand for Prp28 activity using a Prp28 ATPase mutant (Prp28 R499A, a mutation in DEAD-box motif Va) that is proposed to impair ATP hydrolysis and leads to a *cs* growth phenotype (**Fig. 5A**). We predicted that if Luc7 ZnF2_L/L2 places increased demand on Prp28, then this Luc7 mutant should exacerbate the growth defects present with Prp28 R499A. Alternatively, if Luc7 ZnF2_L/L2 facilitates U1/U6 exchange by Prp28 R499A, it should result in suppression of the *cs* phenotype. Luc7 ZnF2_L/L2 not only suppresses the *cs* phenotype, it surpasses Luc7 NΔ31 in its ability to do so (**Fig. 5C**). Together, these data demonstrate that Luc7 ZnF2_L/L2 alters the requirement for ATP hydrolysis by Prp28 during splicing in a way that still requires Prp28 presence and is distinctly different from Luc7 NΔ31. Suppression of Prp28 cold sensitivity is not inconsistent with increasing splicing efficiency at non-consensus splice sites as demonstrated by Luc7 ZnF2_L/L2 in the ACT1-CUP1 reporter assay (**Fig. 3**) or with endogenous 5’ss with U4C substitutions relative to the consensus (**Fig. 4**).

**Figure 5.**
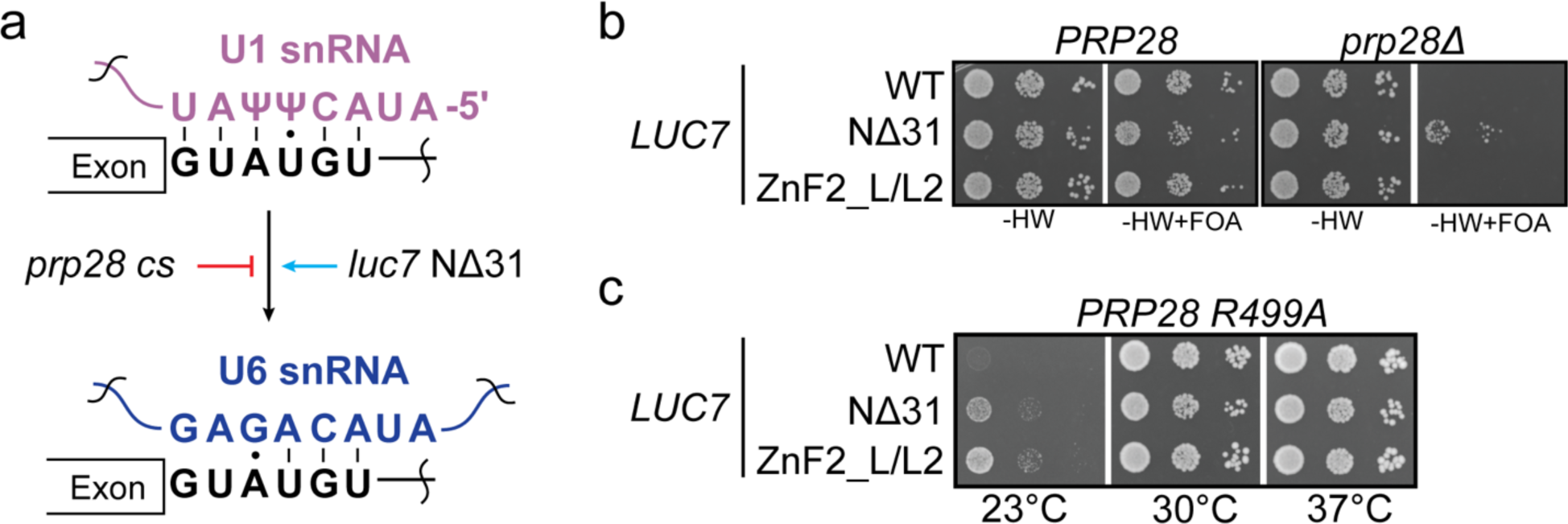
Luc7 ZnF2_L/L2 suppresses the *cs* phenotype of Prp28 R499A. (**a**) Schematic showing the impact of Prp28 alleles that modulate the U1 to U6 exchange during splicing. Truncation of Luc7 bypasses the requirement for Prp28 in splicing and allows splicing to proceed whereas Prp28 ATPase mutants inhibit this step and stall splicing. (**b**) Luc7 ZnF2_L/L2 does not bypass the requirement for Prp28 in splicing unlike Luc7 NΔ31. (**c**) Luc7 ZnF2_L/L2 suppresses cold sensitivity associated with Prp28 R499A.

## DISCUSSION

In this work we studied the roles of the yeast Luc7 ZnF domains in 5’ss selection using humanized variants in which the ZnF domains were mutated to resemble those found in the three human Luc7 paralogs. We were not able to detect an impact of ZnF1 substitutions on splicing of reporter substrates; however, substitutions in ZnF2 that mimic the corresponding domain in human LUC7L and LUC7L2 proteins increased usage of nonconsensus splice sites both in reporter assays and for endogenous yeast transcripts. A mutant with substitutions mimicking ZnF2 of LUC7L3, which is the most sequence divergent of the human paralogs with Luc7, was not viable. In contrast with the Luc7 ZnF2 variant, a Luc7 NΔ31 mutant generally decreased usage of non-consensus 5’ss, including one endogenous cryptic splice site. We also noted that this variant also decreased usage of reporter pre-mRNAs with increased pairing to U1 snRNA (the U4A and U4G ACT1-CUP1 variants) and caused increased accumulation of unspliced transcripts with longer TSS to 5’ss distances. Despite differential impacts on 5’ss usage Luc7 NΔ31 and the ZnF2 variant were both able to suppress a *cs* allele of Prp28, suggesting an impact on both U1 binding to pre-mRNAs and its release during U1/U6 exchange. Altogether, our data reveal how the Luc7 ZnF2 domain can tune 5’ss usage at multiple steps in splicing.

The molecular basis for human LUC7L, LUC7L2, and LUC7L3 promoting different splicing outcomes in human cells is not yet understood. Recently it has been postulated that LUC7L paralog specificity may arise from differences in the number of potential base pairs on either side of the exon/intron junction (Kenny et al. 2022). LUC7L and LUC7L2 (which are more closely related to one another than either is to LUC7L3) may promote splicing primarily at 5’ss with higher levels of complementarity with U1 snRNA at the +3, +4, and +5 positions (CAG/GU<u>AAG</u>U) relative to the −3, −2, and −1 positions (<u>CAG</u>/GUAAGU). These were denoted as having a “right-handed” preference relative to the exon/intron junction (handedness used in this context is unrelated to chirality). In contrast, Luc7L3 appeared to have preference for “left-handed” interactions with higher complementarity on the “exon side” of the junction. Based on the structure of yeast splicing complexes containing Luc7 (**Fig. 1b**, **c**), this would suggest that ZnF2 could play a role in preference for right-handed 5’ss while ZnF1 could play a role in preference for left-handed sites. Yeast 5’ss are much less diverse than their human counterparts with nearly all having extensive complementarity on the intron side (right-handed). Luc7 may function similarly to LUC7L and LUC7L2, and this may explain the lack of effect of ZnF1 changes on reporter and endogenous transcript splicing. In fact, we could find only four yeast genes (∼1% of all pre-mRNAs) with high complementarity within the exon (−3C, −2A, −1G relative to the exon/intron boundary).

Our results with yeast-human Luc7 chimeras also support the notion of a divergent function for LUC7L3. Yeast Luc7 proteins containing human LUC7L3 ZnF1 were viable but those containing LUC7L3 ZnF2 were not, regardless of the origin of ZnF1 (**Fig. 2**). This suggests that the function yeast ZnF2 cannot be compensated for by the more divergent human LUC7L3 ZnF2. For human LUC7L3, this could indicate that while it may promote splicing of left-handed 5’ss, it is also not able to do the same for those that are right-handed. This is in agreement with data from Kenny *et al*. showing that overexpression of LUC7L3 represses splicing of right-handed 5’ss (Kenny et al. 2022).

We were not able to identify a phenotype associated with replacement of yeast ZnF1 with that from any of the human homologs, even when using two different substrates with mutations that strengthened the left-handedness of the 5’ss (**Fig 2b**, **c**). We cannot conclude if this indicates that ZnF1 alone is not responsible for the left-handed preference for 5’ss by LUC7L3 or if this feature of the human splicing machinery cannot be recapitulated in our yeast system. Nonetheless, it suggests that further work is needed to define the role of ZnF1. Work from Agarwal *et al*. indicated that while yeast were viable with Luc7 containing a disrupted metal-binding site in ZnF1, several of these mutants possessed a *ts* phenotype and were synthetic lethal with mutations in a variety of other splicing factors (Agarwal et al. 2016). This indicates that ZnF1 does have a function in yeast splicing, but its role may be context-dependent.

One of the complexities in studying U1 snRNP-associated factors *in vivo* is that splicing outcomes (for example, as measured by RNA-seq) are only correlated with U1 association. For splicing chemistry to occur, it is essential that U1 be released during the transition from the pre-B to B complex spliceosome. This release must occur so that 5’ss interactions with the U1 snRNA can be exchanged for those with the U6 snRNA. This normally requires the action of the DEAD-box ATPase Prp28. Consequently, inferring changes in U1 binding from measurements of mRNA production are difficult and a convolution of U1 overall binding properties (on- and off-rates), U1 release by Prp28, and U6 binding among likely other influences. Our work shows that while substitution of ZnF2 of Luc7 can result in relatively few overall changes in the transcriptome (**Fig. 4**), it can still perturb the ATPase-dependence of U1 release. Humanization of Luc7 ZnF2 does not bypass the need for Prp28 but does suppress the *cs* phenotype of a Prp28 ATPase site mutant (**Fig. 5**). This suggests that human LUC7L/L2 ZnF2 facilitates U1 release by Prp28. We do not know if this occurs by weakening the U1 snRNA/5’ss duplex at this stage and/or by stimulating Prp28 activity. However, it does provide additional evidence that the zinc fingers play roles in both recognition of the 5’ss by U1 and its release during activation. Given the conservation of these factors, it is likely that this is also true in humans. Thus, understanding the function of the human Luc7 homologs should consider their impact on both U1 binding and release. Given that LUC7L2 promotes glycolysis at the expense of oxidative phosphorylation (Jourdain et al. 2021), it is tempting to speculate that different Luc7 homologs have different ATPase dependencies for the U1/U6 exchange that could correlate with cellular metabolic state.

## MATERIALS AND METHODS

*Saccharomyces cerevisiae* strains used in these studies were derived from 46α (kind gift of David Brow) or a *PRP28* and *LUC7* double shuffle strain (gift of Beate Schwer) (Lesser and Guthrie 1993a; Agarwal et al. 2016). The open reading frame of *LUC7* and the 500 base pairs up and downstream was amplified from yeast genomic DNA and cloned into the CEN6/ARS4 centromeric plasmids pRS416 and pRS414. **Supplemental Tables 1 and 2** contain detailed lists of strains and plasmids used. Yeast transformation and growth was carried out using standard techniques and media (Amberg et al. 2005).

### Site-Directed Mutagenesis

Point mutants were generated using inverse polymerase chain reaction (PCR) with Phusion DNA polymerase (New England Biolabs) or Herculase II (Agilent). PCR was performed for 16 cycles using primers with the desired nucleotides changes incorporated at or near the 5’ ends. Template DNA was removed by treatment with DpnI (New England Biolabs), and the PCR products were subsequently 5’ phosphorylated using T4 polynucleotide kinase (New England Biolabs) and self-ligated by T4 DNA ligase (New England Biolabs) before being used to transform Top10 competent cells (Thermo Fisher Scientific). Individual colonies were screened by Sanger sequencing to identify the desired changes.

### Temperature Growth Assays

Yeast strains were grown to mid-log phase in YPD supplemented with 0.003% w/v adenine hemisulfate (YPAD) or selective media, the OD_600_ was adjusted to 0.5 and equal volumes were spotted onto YPAD or selective plates. Plates were incubated at the indicated temperature and scored after 3 days growth at 30°C or 3 days growth at 23°C or 37°C.

### *ACT1-CUP1* Copper Assays

*ACT1-CUP1* reporters and growth assays have been described previously (Lesser and Guthrie 1993a; Carrocci et al. 2017). Briefly, yeast strains expressing WT or mutant proteins and *ACT1-CUP1* reporters were grown to mid-log phase in SC-LEU media to maintain selection for the reporter plasmids, adjusted to OD_600_ = 0.5 and equal volumes were spotted onto plates containing 0, 0.025, 0.05, 0.075, 0.1, 0.15, 0.2, 0.25, 0.3, 0.4, 0.5, 0.6, 0.7, 0.8, 0.9, 1.0, 1.1, 1.2, 1.3, 1.4, 1.5, 1.6, 1.7, 1.8, 1.9, 2.0, 2.25, 2.5, 3.0, or 3.5 mM CuSO_4_. Plates were scored after 3 days growth at 30°C.

### RT-PCR RNA Analysis

Yeast were grown in YPAD media until OD_600_ reached 0.5–0.8. Cells (8 OD_600_ units) were harvested by centrifugation, washed with water and total cellular RNA was isolated using a MasterPure Yeast RNA Purification Kit (Lucigen) according to the vendor’s instructions. DNaseI-treated total RNA (4 µg) was reverse transcribed using Primescript reverse transcriptase (Takara Bio) and random hexamers (Thermo Fisher Scientific). Assembled reactions were incubated at 30°C for 10 min, followed by 1 h at 42°C for complete extension. The RT was heat inactivated by treatment at 70°C for 15 min and reactions were diluted 1:20 and used in PCR without further purification. PCR reactions were carried out using Taq polymerase (New England Biolabs) and contained 2.5 µM gene-specific primers that flanked the intron. One of the primers was also labeled at the 5’ end with Cy5 to facilitate fluorescence imaging. Products were separated using 2.5% (w/v) metaphor agarose (Lonza) in 1×TBE, and the gel was subsequently imaged using an iBright imager (Thermo Fisher Scientific). Band intensities were quantified using ImageJ.

### RNA extraction and RNA Sequencing

RNA was isolated as previously described (Wang et al. 2020). Briefly, strains expressing either *LUC7-WT* or mutants *luc7-nΔ31* and *luc7-znf2* and with *UPF1* deleted were grown overnight in YPD at 30°C. The following morning, the cultures were diluted in fresh YPD to OD_600_ = 0.1 and grown at 30°C for 2 doublings; after which cells were collected and flash frozen in liquid nitrogen. Total RNA was extracted using hot phenol:chloroform followed by ethanol precipitation. Total RNA (40 µg) from each condition was then treated with DNaseI (Invitrogen) to remove genomic DNA. Samples were ribo-depleted using the Ribocop for Yeast kit (Lexogen) and libraries were prepared using the CORALL Total RNA-Seq V2 kit (Lexogen). Barcoding was carried out using the UDI 12 nt Set B1 (Lexogen), and amplification was done using 15 cycles of PCR. Sequencing was performed on a NovaSeq PE150 by Novogene who also carried out sample demultiplexing.

### Read Quality Control and Mapping

UMI-tools was used to extract unique molecular identifiers (UMI) from reads (Smith et al. 2017). Each UMI was 12 nucleotides (nt) long (--bc-pattern=NNNNNNNNNNNN). Cutadapt was used for quality control and to remove adaptors and polyA sequences (Martin 2011). Terminal “N’’s were removed from each read. Adapters-g “T{100}”-a AGATCGGAAGAGC-A AGATCGGAAGAGC-A “A{100}”. A minimum overlap of 2 nt was required for each adapter and a maximum of 2 errors was allowed in each adapter sequence. Bases with quality scores less than 20 were removed from both ends of each read. A minimum read length of 50 nt was required. Any read with more than 4 ambiguous bases (N) after filtering was removed. Reads were aligned to the *Saccharomyces* genome (S288C version R64-3-1) downloaded using HISAT2 (version: 2.2.1) (Cherry et al. 2012; Engel et al. 2014; Kim et al. 2019). Known splice site annotations were acquired from the SGD. For non-canonical splicing events a minimum intron length of 20 nt and a maximum intron length of 1,000 nt was required. A penalty of 0 was set for non-canonical splice site alignment. PCR duplicates were removed using umi_tools extract prior to counting.

### Counting of mRNAs, Spliced, and Unspliced Reads

Genome annotation files were acquired from the SGD. Counting was done using Rsubread featureCounts (Liao et al. 2019). Each read in a pair was counted individually and multiple overlapping was allowed. DESeq2 was used to estimate differential gene expression (Love et al. 2014). To count unambiguously unspliced reads, both sides of each intron were counted separately, and the results were averaged. The annotation file was 1 nt outside of the intron and 1 nt inside the intron; a read needed to be mapped to both positions to be counted. Only non-split reads were counted, and each read in a pair was counted individually with multiple overlapping allowed. To count unambiguously spliced reads, both sides of each intron were counted separately, and the results were averaged. The annotated file was 5 nt outside of the intron and a minimum overlap of 4 nt was required. Only split alignments were counted. In some cases, alternative splice sites were counted as spliced reads; however, this was a small minority of reads. Each read in a pair was counted individually and multiple overlapping was allowed. Any introns with no spliced reads in any replicate were removed from further analysis. The implementation can be found in the counting script (https://github.com/SamDeMario-lab/Luc7_splicing).

### Splicing Efficiency Calculations

Splicing efficiency was calculated as the ratio of reads which were unambiguously unspliced over unambiguously spliced for each intron. This ratio was calculated individually for each replicate and the geometric mean was used as the splicing efficiency for downstream analysis. We note that due to PCR biases our splicing efficiency calculations are unlikely to be accurate representations of the true levels of spliced and unspliced transcripts. However, all of our analysis is based around relative changes in splicing efficiency and, therefore, the biases are uniform across all samples. The “Intronic reads in mutant/WT” ratio was calculated as the splicing efficiency of the mutant divided by the splicing efficiency of from strains expressing WT Luc7.

### Data Processing and Plot Generation

Data was processed and visualized in R version 4.2.2 (R Core Development Team 2022). DESeq2 was used to test differential expression (Love et al. 2014). Plots were generated using ggplot2 (Wickham 2016), ggforce was used to create reversed log transformed axes (Pedersen 2022), and ggseqlogo was used to make sequence logos (Wagih 2017). Rcolorbrewer was used to select color pallets for most panels (Neuwirth 2022). Figures were arranged using gridExtra (Auguie 2017). BSgenome.Scerevisiae.UCSC.sacCer3, biostrings and, biomartr were used to extract reference sequences from genomic coordinates (The Bioconductor Dev Team 2014; Drost and Paszkowski 2017; Pagès et al. 2022). Likely branch point sequences were acquired from the Ares lab Yeast Intron Database (Grate 2002). Any intron without a predicted branchpoint was excluded from the branch point analysis.

## ACKNOWLEDGEMENTS

We thank members of the Hoskins lab for their helpful discussions. We thank Dave Brow, Charles Query, and Beate Schwer for strains and plasmids used in this study.

## DATA AND CODE AVAILABILITY

All scripts used to process data as well as count tables and annotation files are available on GitHub (https://github.com/SamDeMario-lab/Luc7_splicing). All data generated during this study are available on the GEO at accession number (PRJNA1022512).

## REFERENCES

Agarwal R, Schwer B, Shuman S. 2016. Structure-function analysis and genetic interactions of the Luc7 subunit of the Saccharomyces cerevisiae U1 snRNP. RNA 22: 1302–1310.

Amberg DC, Burke D, Strathern. JN. 2005. Methods in Yeast Genetics: A Cold Spring Harbor Laboratory Course Manual. Cold Spring Harbor Laboratory Press, Cold Spring Harbor, NY.

Auguie B. 2017. gridExtra: Miscellaneous Functions for “Grid” Graphics.

Carrocci TJ, Zoerner DM, Paulson JC, Hoskins AA. 2017. SF3b1 mutations associated with myelodysplastic syndromes alter the fidelity of branchsite selection in yeast. Nucleic Acids Res 45: 4837–4852.

Chalivendra S, Shi S, Li X, Kuang Z, Giovinazzo J, Zhang L, Rossi J, Saviola AJ, Wang J, Welty R et al. 2023. Selected humanization of yeast U1 snRNP leads to global suppression of pre-mRNA splicing and mitochondrial dysfunction in the budding yeast. bioRxiv. doi:10.1101/2023.12.15.571893.

Cherry JM, Hong EL, Amundsen C, Balakrishnan R, Binkley G, Chan ET, Christie KR, Costanzo MC, Dwight SS, Engel SR et al. 2012. Saccharomyces Genome Database: the genomics resource of budding yeast. Nucleic Acids Res 40: D700–705.

Daniels NJ, Hershberger CE, Gu X, Schueger C, DiPasquale WM, Brick J, Saunthararajah Y, Maciejewski JP, Padgett RA. 2021. Functional analyses of human LUC7-like proteins involved in splicing regulation and myeloid neoplasms. Cell Rep 35: 108989.

Drost HG, Paszkowski J. 2017. Biomartr: genomic data retrieval with R. Bioinformatics 33: 1216–1217.

Eaton SL, Roche SL, Llavero Hurtado M, Oldknow KJ, Farquharson C, Gillingwater TH, Wishart TM. 2013. Total protein analysis as a reliable loading control for quantitative fluorescent Western blotting. PLoS One 8: e72457.

Engel SR, Dietrich FS, Fisk DG, Binkley G, Balakrishnan R, Costanzo MC, Dwight SS, Hitz BC, Karra K, Nash RS et al. 2014. The reference genome sequence of Saccharomyces cerevisiae: then and now. G3 (Bethesda) 4: 389-398.

Espinosa S, De Bortoli F, Li X, Rossi J, Wagley ME, Lo HG, Taliaferro JM, Zhao R. 2022. Human PRPF39 is an alternative splicing factor recruiting U1 snRNP to weak 5’ splice sites. RNA 29: 97–110.

Fica SM. 2020. Cryo-EM snapshots of the human spliceosome reveal structural adaptions for splicing regulation. Curr Opin Struct Biol 65: 139–148.

Fortes P, Bilbao-Cortes D, Fornerod M, Rigaut G, Raymond W, Seraphin B, Mattaj IW. 1999. Luc7p, a novel yeast U1 snRNP protein with a role in 5’ splice site recognition. Genes Dev 13: 2425–2438.

Gornemann J, Kotovic KM, Hujer K, Neugebauer KM. 2005. Cotranscriptional spliceosome assembly occurs in a stepwise fashion and requires the cap binding complex. Mol Cell 19: 53–63.

Grate L, Ares M. 2002. Searching Yeast Intron Data at the Areslab Website. (in Guide to Yeast Genetics and Molecular and Cell Biology, Part B, C. Guthrie and G. Fink, eds) Methods Enz. 350: 380–392.

Hackmann A, Wu H, Schneider UM, Meyer K, Jung K, Krebber H. 2014. Quality control of spliced mRNAs requires the shuttling SR proteins Gbp2 and Hrb1. Nat Commun 5: 3123.

Hansen SR, White DS, Scalf M, Correa IR, Smith LM, Hoskins AA. 2022. Multi-step recognition of potential 5’ splice sites by the Saccharomyces cerevisiae U1 snRNP. Elife 11.

Jourdain AA, Begg BE, Mick E, Shah H, Calvo SE, Skinner OS, Sharma R, Blue SM, Yeo GW, Burge CB et al. 2021. Loss of LUC7L2 and U1 snRNP subunits shifts energy metabolism from glycolysis to OXPHOS. Mol Cell 81: 1905–1919 e1912.

Kandels-Lewis S, Seraphin B. 1993. Involvement of U6 snRNA in 5’ splice site selection. Science 262: 2035–2039.

Kawashima T, Douglass S, Gabunilas J, Pellegrini M, Chanfreau GF. 2014. Widespread use of non-productive alternative splice sites in Saccharomyces cerevisiae. PLoS Genet 10: e1004249.

Kenny CJ, McGurk MP, Burge CB. 2022. Human LUC7 proteins impact splicing of two major subclasses of 5’ splice sites. Biorxiv. doi:10.1101/2022.12.07.519539.

Kim D, Paggi JM, Park C, Bennett C, Salzberg SL. 2019. Graph-based genome alignment and genotyping with HISAT2 and HISAT-genotype. Nat Biotechnol 37: 907–915.

Kondo Y, Oubridge C, van Roon AM, Nagai K. 2015. Crystal structure of human U1 snRNP, a small nuclear ribonucleoprotein particle, reveals the mechanism of 5’ splice site recognition. Elife 4.

Larson JD, Hoskins AA. 2017. Dynamics and consequences of spliceosome E complex formation. Elife 6.

Lesser CF, Guthrie C. 1993a. Mutational analysis of pre-mRNA splicing in Saccharomyces cerevisiae using a sensitive new reporter gene, CUP1. Genetics 133: 851-863.

-. 1993b. Mutations in U6 snRNA that alter splice site specificity: implications for the active site. Science 262: 1982–1988.

Li X, Liu S, Jiang J, Zhang L, Espinosa S, Hill RC, Hansen KC, Zhou ZH, Zhao R. 2017. CryoEM structure of Saccharomyces cerevisiae U1 snRNP offers insight into alternative splicing. Nat Commun 8: 1035.

Li X, Liu S, Zhang L, Issaian A, Hill RC, Espinosa S, Shi S, Cui Y, Kappel K, Das R et al. 2019. A unified mechanism for intron and exon definition and back-splicing. Nature 573: 375–380.

Liao Y, Smyth GK, Shi W. 2019. The R package Rsubread is easier, faster, cheaper and better for alignment and quantification of RNA sequencing reads. Nucleic Acids Res 47: e47.

Love MI, Huber W, Anders S. 2014. Moderated estimation of fold change and dispersion for RNA-seq data with DESeq2. Genome Biol 15: 550.

Love SL, Emerson JD, Koide K, Hoskins AA. 2023. Pre-mRNA splicing-associated diseases and therapies. RNA Biol 20: 525–538.

Martin M. 2011. Cutadapt removes adapter sequences from high-throughput sequencing reads. EMBnetjournal 17.

Matlin AJ, Clark F, Smith CW. 2005. Understanding alternative splicing: towards a cellular code. Nat Rev Mol Cell Biol 6: 386–398.

Neuwirth E. 2022. RColorBrewer: ColorBrewer Palettes.

Pagès H, Aboyoun P, Gentleman R, DebRoy S. 2022. Biostrings: Efficient manipulation of biological strings.

Pedersen TL. 2022. ggforce: Accelerating ‘ggplot2’.

Plaschka C, Lin PC, Charenton C, Nagai K. 2018. Prespliceosome structure provides insights into spliceosome assembly and regulation. Nature 559: 419–422.

Puig O, Bragado-Nilsson E, Koski T, Seraphin B. 2007. The U1 snRNP-associated factor Luc7p affects 5’ splice site selection in yeast and human. Nucleic Acids Res 35: 5874–5885.

R Core Development Team. 2022. R: A language and environment for statistical computing. R Foundation for Statistical Computing.

Roca X, Krainer AR, Eperon IC. 2013. Pick one, but be quick: 5’ splice sites and the problems of too many choices. Genes Dev 27: 129–144.

Ruby SW, Abelson J. 1988. An early hierarchic role of U1 small nuclear ribonucleoprotein in spliceosome assembly. Science 242: 1028–1035.

Sayani S, Janis M, Lee CY, Toesca I, Chanfreau GF. 2008. Widespread impact of nonsense-mediated mRNA decay on the yeast intronome. Mol Cell 31: 360–370.

Schwer B, Shuman S. 2014. Structure-function analysis of the Yhc1 subunit of yeast U1 snRNP and genetic interactions of Yhc1 with Mud2, Nam8, Mud1, Tgs1, U1 snRNA, SmD3 and Prp28. Nucleic Acids Res 42: 4697-4711.

Shcherbakova I, Hoskins AA, Friedman LJ, Serebrov V, Correa IR, Jr., Xu MQ, Gelles J, Moore MJ. 2013. Alternative spliceosome assembly pathways revealed by single-molecule fluorescence microscopy. Cell Rep 5: 151–165.

Sievers F, Higgins DG. 2014. Clustal Omega, accurate alignment of very large numbers of sequences. Methods Mol Biol 1079: 105–116.

Smith T, Heger A, Sudbery I. 2017. UMI-tools: modeling sequencing errors in Unique Molecular Identifiers to improve quantification accuracy. Genome Res 27: 491–499.

Spingola M, Grate L, Haussler D, Ares M, Jr. 1999. Genome-wide bioinformatic and molecular analysis of introns in Saccharomyces cerevisiae. RNA 5: 221–234.

Staley JP, Guthrie C. 1999. An RNA switch at the 5’ splice site requires ATP and the DEAD box protein Prp28p. Mol Cell 3: 55–64.

The Bioconductor Dev Team. 2014. BSgenome.Scerevisiae.UCSC.sacCer3: Saccharomyces cerevisiae (Yeast) full genome (UCSC version sacCer3).

Wagih O. 2017. ggseqlogo: a versatile R package for drawing sequence logos. Bioinformatics 33: 3645–3647.

Wang C, Liu Y, DeMario SM, Mandric I, Gonzalez-Figueroa C, Chanfreau GF. 2020. Rrp6 Moonlights in an RNA Exosome-Independent Manner to Promote Cell Survival and Gene Expression during Stress. Cell Rep 31: 107754.

Wickham H. 2016. ggplot2: Elegant Graphics for Data Analysis. Springer-Verlag, New York.

Wilkinson ME, Charenton C, Nagai K. 2020. RNA Splicing by the Spliceosome. Annu Rev Biochem 89: 359-388.

Zhang X, Zhan X, Bian T, Yang F, Li P, Lu Y, Xing Z, Fan R, Zhang QC, Shi Y. 2024. Structural insights into branch site proofreading by human spliceosome. Nat Struct Mol Biol.

