## Supplemental Figures for "Functional Analysis of the Zinc Finger Modules of the *S. cerevisiae* Splicing Factor Luc7"

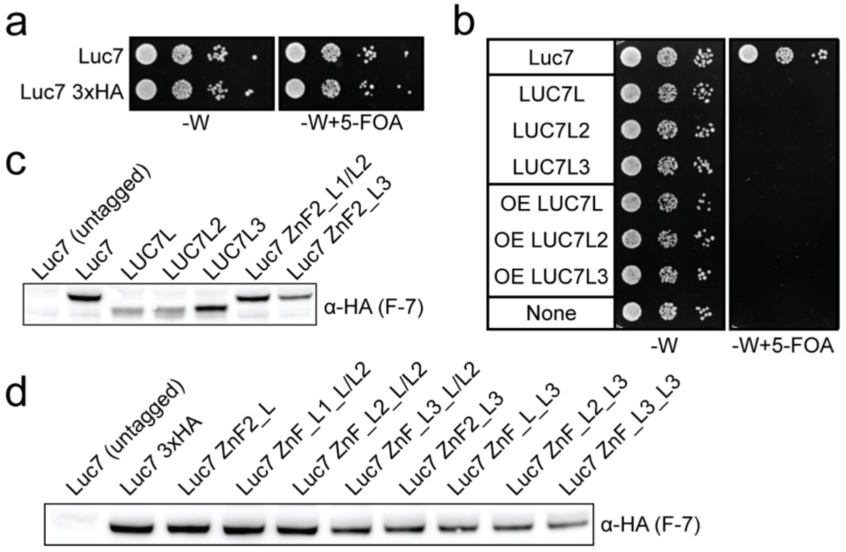


**Figure S1. Expression of mutant Luc7 in yeast.** (**a**) C-terminal HA-tagging of Luc7 has no effect on yeast growth compared to the untagged control. (**b**) Human LUC7L proteins (truncated before the RS domain) cannot support growth in yeast as the sole copy of *LUC7*. Overexpression (OE) of the proteins from a *TDH3* promoter does not rescue growth. (**c**) Western blotting using the F-7 anti-HA tag antibody shows expression LUC7L, LUC7L2, LUC7L3 and ZnF2 mutants before FOA shuffle in merodiploid strains. (**d**) Expression levels of ZnF mutant Luc7 proteins in yeast. For these blots, yeast extracts were normalized for total protein content as a loading control and equivalent total amounts of protein were added to each well prior to SDS-PAGE (Eaton et al. 2013).

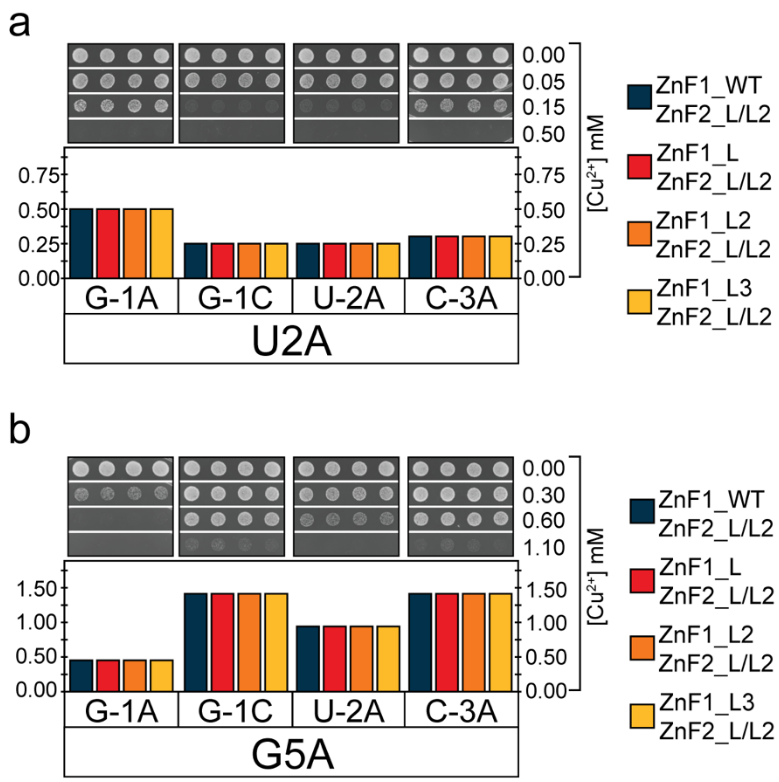


**Figure S2. Modifying ZnF1 and ZnF2 simultaneously does not modify 5'ss usage.** (**a**) Growth on $Cu^{2+}$-containing media of yeast strains with reporters containing substitutions of the U+2 position in addition to the indicated changes at the -1 to -3 sites are unaffected. (**b**) Same as in (a) except with a substitution at G+5. Shown is a representative ACT1-CUP1 assay from three experimental replicates.

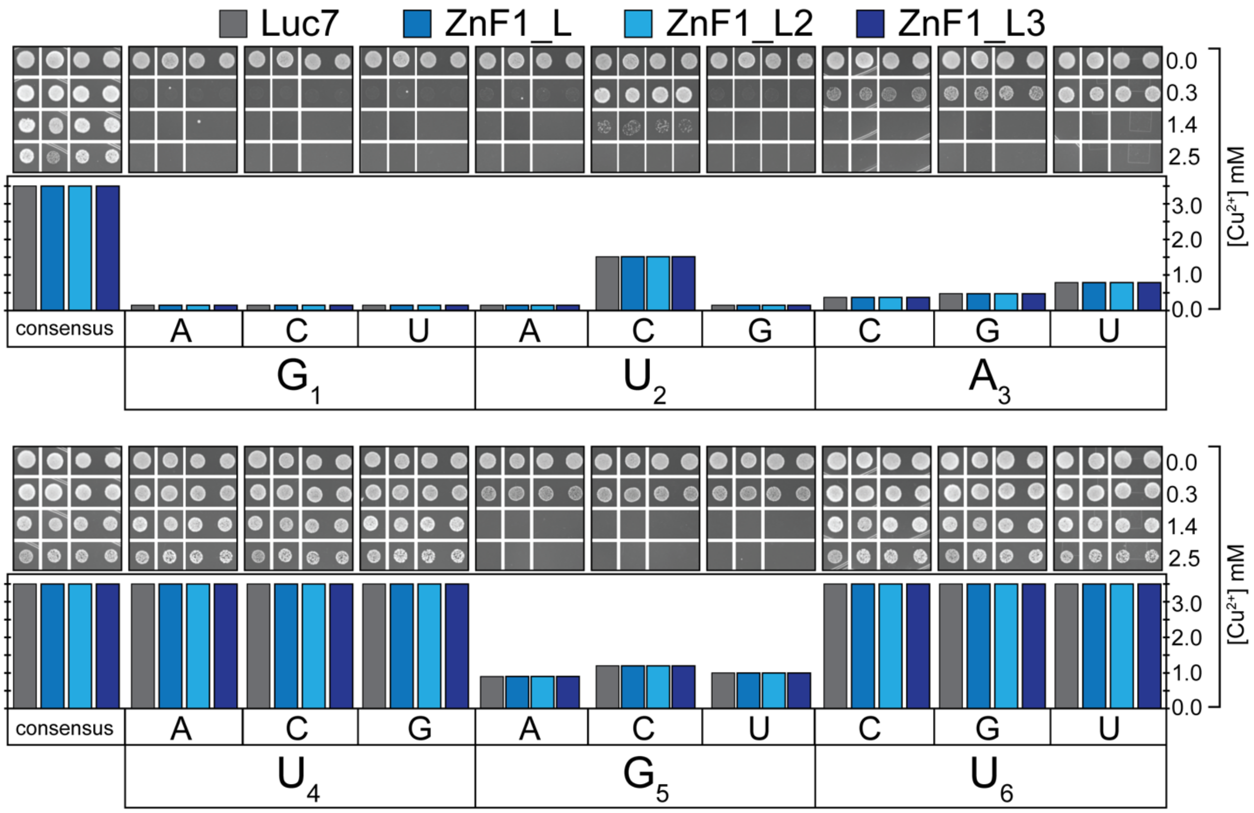


**Figure S3. Luc7 ZnF1 mutants do not impact non-consensus 5'ss usage**. *ACT1-CUP1* reporters with single substitutions at all positions of the 5'ss have similar $Cu^{2+}$ tolerances with all ZnF1 mutants tested. Shown is a representative ACT1-CUP1 assay from three experimental replicates.

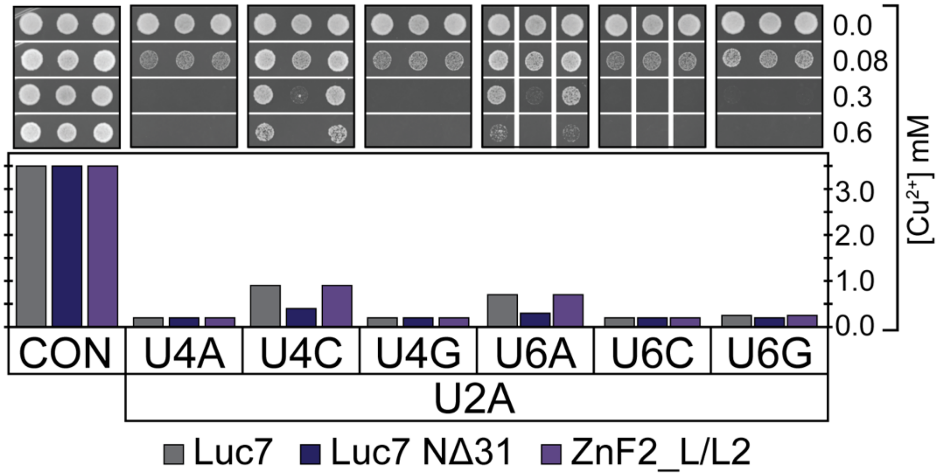


**Figure S4. ACT1-CUP1 assays using reporters that combine U2A with substitutions at positions U4 and U6.** Shown is a representative ACT1-CUP1 assay from three experimental replicates.

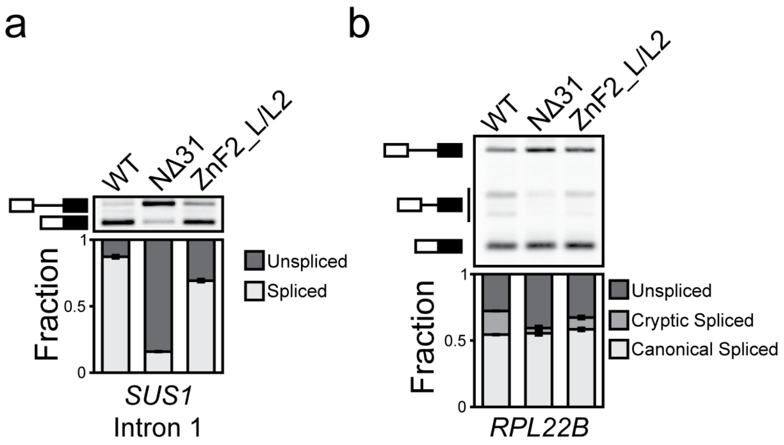


**Figure S5. RT-PCR analysis of splicing changes in endogenous transcripts upon mutation of Luc7.** (**a**) Luc7 ZnF2_L/L2 or NΔ31 accumulate unspliced pre-mRNA from the first intron of *SUS1*. (**b**) *RPL22B* unspliced pre-mRNA accumulates in Luc7 mutants but less cryptic splicing is observed with NΔ31. Error bars represent standard deviation from three independent replicates.

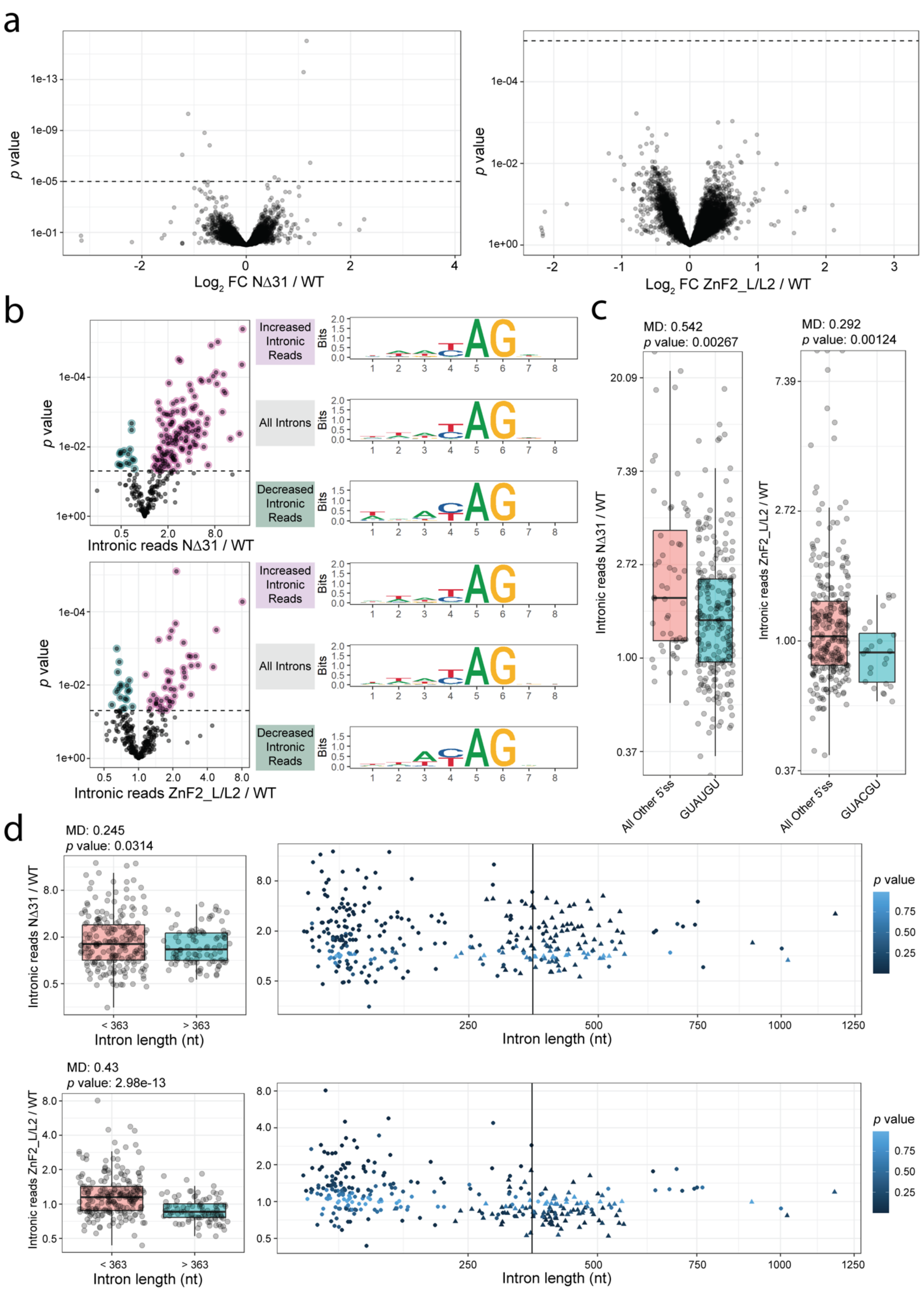
a
p value
Log2 FC NΔ31 / WT
Log2 FC ZnF2_L/L2 / WT
b
Increased Intronic Reads
All Introns
Decreased Intronic Reads
Bits
Intronic reads NΔ31 / WT
Intronic reads ZnF2_L/L2 / WT
c
MD: 0.542
p value: 0.00267
MD: 0.292
p value: 0.00124
All Other 5'ss
GUAUGU
GUACGU
d
MD: 0.245
p value: 0.0314
MD: 0.43
p value: 2.98e-13
Intron length (nt)
< 363
> 363

**Figure S6. Supplemental analysis of RNA-seq data from cells expressing WT and mutant forms of Luc7.** (**a**) mRNA levels for WT and Luc7 variant strains. mRNA annotation from SGD. Horizontal line indicates *p* value = 1e-5. (**b**) Plots showing change in splicing efficiency for each intron in Luc7 NΔ31 and Luc7 ZnF2_L/L2. Sequence logos of 3'ss and surrounding bases. (**c**) Unequal variances t-tests using geometric means were performed for each mutant testing the common 5'ss in yeast ("GUAUGU", "GUAAGU", "GUACGU", "GUAUGA", "GUAUGC") against all other 5'ss. Only 5'ss with *p* value < 0.05 were reported. (**d**) Splicing efficiency vs intron length for Luc7 NΔ31 (Top) and Luc7 ZnF2_L/L2 (Bottom). Intron length threshold was determined by minimizing *p* values for Luc7 ZnF2_L/L2 along a sliding window of intron lengths between 50 and 500 nt. Data point shape indicates ribosomal protein genes (triangle) or non-ribosomal protein genes (circle). Color indicates *p* value as in Figure 4C. (MD - Difference of geometric means)
